# Efficient linear dsDNA tagging using deoxyuridine excision

**DOI:** 10.1101/2021.06.29.450395

**Authors:** Eric J. Strobel

## Abstract

Site-specific strategies for exchanging segments of dsDNA are important for DNA library construction and molecular tagging. Deoxyuridine (dU) excision is an approach for generating 3’ ssDNA overhangs in gene assembly and molecular cloning procedures. Unlike approaches that use a multi-base pair motif to specify a DNA cut site, dU excision requires only a dT→dU substitution. Consequently, excision sites can be embedded in biologically active DNA sequences by placing dU substitutions at non-perturbative positions. In this work, I describe a molecular tagging method that uses dU excision to exchange a segment of a dsDNA strand with a long synthetic oligonucleotide. The core workflow of this method, called <u>d</u>eoxy<u>U</u>ridine e<u>X</u>cision-tagging (dUX-tagging), is an efficient one-pot reaction: strategically positioned dU nucleotides are excised from dsDNA to generate a 3’ overhang so that additional sequence can be appended by annealing and ligating a tagging oligonucleotide. The tagged DNA is then processed by one of two procedures to fill the 5’ overhang and remove excess tagging oligo. To facilitate its widespread use, all dUX-tagging procedures exclusively use commercially available reagents. As a result, dUX-tagging is a concise and easily implemented approach for high-efficiency linear dsDNA tagging.

## Introduction

Molecular barcoding is a strategy for tagging nucleic acids with sequence-encoded identifiers to distinguish between multiple samples or individual molecules (1–3). In genomics applications, molecular tags are typically appended using procedures that are not site-specific in order to accommodate variable nucleic acid ends. However, in high-throughput experiments that rely on synthetic DNA libraries with defined ends, site-specific methods for appending or exchanging DNA sequences concisely and with high efficiency are desirable.

Deoxyuridine (dU) excision is an established strategy for generating 3’ overhangs during molecular cloning and gene assembly procedures (4–9). In this approach, dU nucleotides are excised from DNA through two enzymatic reactions: After the uracil base is excised by a uracil DNA glycosylase (10,11), the resulting abasic site is excised by an AP lyase such as endonuclease III (12,13) or endonuclease VIII (14,15). These reactions, which are readily performed together using a commercially available <u>U</u>racil-<u>S</u>pecific <u>E</u>xcision <u>R</u>eagent (USER^®^) enzyme mixture (New England Biolabs), generate a one nucleotide gap with a 5’ phosphate that can be used to join DNA molecules. A key advantage of dU excision-based cloning methods is that the 3’ overhang used for DNA end-joining is specified by a dT→dU substitution rather than the presence of a more complex recognition sequence. Consequently, dU excision is an ideal strategy for encoding molecular tagging functionality in biologically active DNA sequences and constant regions of complex DNA libraries.

In this work, I describe a site-specific, sequence-independent method for tagging dsDNA with a molecular barcode in a one-pot reaction. In this procedure, called <u>d</u>eoxy<u>U</u>ridine e<u>X</u>cision-tagging (dUX-tagging), strategically positioned dU nucleotides are excised to generate a long 3’ overhang so that a synthetic ‘tagging oligonucleotide’ that contains a molecular barcode can be annealed and ligated to the target DNA molecule, leaving a 5’ overhang. I describe two procedures for removing excess tagging oligo and filling the 5’ overhang that yield tagged dsDNA without the formation of any side products. Because tagging sites are encoded by dU nucleotides rather than a recognition motif with a defined sequence, dUX-tagging is compatible with complex DNA sequence libraries. This sequence-independence also enables dUX-tagging sites to be integrated with functional DNA elements, such as the *E*. *coli* σ^70^ promoter used for method development here. Overall, this work establishes dUX-tagging as a flexible and lightweight strategy for embedding molecular tagging functionality in arbitrary DNA sequences.

## Materials and Methods

### Oligonucleotides

All oligonucleotides were purchased from Integrated DNA Technologies (IDT). A detailed description of all oligonucleotides including sequence, modifications, and purifications is presented in Supplementary Table S1.

### Proteins

All proteins, including Q5^®^ High-Fidelity DNA Polymerase, Q5U^®^ High-Fidelity DNA Polymerase, Vent^®^ (exo-) DNA polymerase, Vent^®^ DNA polymerase, *Sulfolobus* DNA Polymerase IV, Thermolabile Exonuclease I, Klenow Fragment (3’→5’ exo-), Klenow Fragment, T4 DNA Polymerase, Thermolabile USER^®^ II Enzyme, T4 DNA Ligase, *E*. *coli* RNA Polymerase holoenzyme, ET SSB, Thermolabile Proteinase K, and RNase I_f_ were purchased from New England Biolabs (NEB).

### DNA template preparation

DNA templates were prepared exactly as described previously (16). Supplementary Table S2 provides details on the oligonucleotides and specific processing steps used for every DNA template preparation in this work.

### Sequences

Annotated sequences of the substrate DNA template (https://benchling.com/s/seq-L8QDkyWpnMGkdqnkOXlO) and fully tagged DNA template (https://benchling.com/s/seq-i86HHdmB4YQdE90bVegJ) are available at Benchling; Unannotated sequences are shown in Supplementary Table S3. The Illumina adapter sequences used in these constructs are from the TrueSeq Small RNA kit. In the sequence annotations, VRA3 is the reverse complement of the Illumina RA5 adapter, and VRA5 is the reverse complement of the Illumina RA3 adapter; this notation was originally used in the Precision Run-On Sequencing method (17,18).

### Preparation of streptavidin-coated magnetic beads

10 μl of Dynabeads™ MyOne™ Streptavidin C1 beads (Invitrogen) per 25 μl sample volume were prepared in bulk essentially as described previously (16). Briefly, storage buffer was removed and the beads were resuspended in 500 μl of Hydrolysis Buffer (100 mM NaOH, 50 mM NaCl) and incubated at room temperature for 10 minutes with rotation. Hydrolysis Buffer was removed, and the beads were resuspended in 1 ml of High Salt Wash Buffer (50 mM Tris-HCl (pH 7.5), 2 M NaCl, 0.5% Triton X-100), transferred to a new tube, and washed by rotating for 5 minutes at room temperature. High Salt Wash Buffer was removed, and the beads were resuspended in 1 ml of Binding Buffer (10 mM Tris-HCl (pH 7.5), 300 mM NaCl, 0.1 % Triton X-100), transferred to a new tube, and washed by rotating for 5 minutes at room temperature. Binding buffer was removed, and the beads were resuspended in 25 of Binding Buffer of per sample and transferred to a new tube. Beads were prepared fresh for each experiment and kept on ice until use.

### Deoxyuridine excision-tagging (dUX-tagging)

dUX-tagging reactions were performed using a master mix from which fractions were taken at various points during processing to visualize intermediate steps. The starting volume of an individual sample within the master mix was 25 μl. For clarity and simplicity, the procedure below specifies the volume of each reagent added to the master mix per 25 μl sample volume at each step. A description of the intermediate fractions that were collected is provided at the end of this sub-section.

#### USER digestion, tagging oligonucleotide annealing, and DNA ligation

A reaction master mix containing 1X T4 DNA Ligase Buffer (NEB) (2.5 μl/sample volume of 10X T4 DNA Ligase Buffer), 5 nM DNA template (0.125 pmol/sample volume), and 0.02 U/μl Thermolabile USER^®^ II Enzyme (NEB) (0.5 μl/sample volume) was prepared on ice in a thin-walled 200 μl tube. The dU excision reaction was incubated at 37 °C for 30 min on a thermal cycler block with a heated lid set to 45 °C. After the 30 min incubation the reaction temperature was held at 8 °C. After dU excision, 0.5 μl/sample volume of 2.5 μM tagging oligo VRA5_16N_PRA1m12 or VRA5_16N_3T_PRA1m12 (Figure 6C,D and Supplementary Figure S2 only) (sequences are available in Supplementary Table S1) was added to the master mix. To anneal the tagging oligo and inactivate Thermolabile USER^®^ II Enzyme, the master mix was placed on a thermal cycler block set to 70 °C with a heated lid set to 105 °C and slowly cooled using the protocol: 70 °C for 5 minutes, ramp to 65 °C at 0.1 °C/s, 65 °C for 5 minutes, ramp to 60 °C at 0.1 °C/s, 60 °C for 2 minutes, ramp to 25 °C at 0.1 °C/s, hold at 25 °C. After annealing the tagging oligo, 1 μl/sample volume of T4 DNA ligase (NEB) was added to the master mix. The ligation reaction was incubated at 25 °C for 1 hour.

#### Bead-based tagging oligonucleotide clean-up and low-temperature primer extension

After the ligation step was completed, T4 DNA ligase was inactivated by incubation at 65 °C for 10 minutes. The master mix was transferred to a 1.7 ml microcentrifuge tube and mixed with 26.5 μl/sample volume of 2X Binding Buffer (20 mM Tris-HCl, (pH 7.5), 600 mM NaCl, 0.2% Triton X-100). Pre-equilibrated streptavidin-coated magnetic beads were placed on a magnet stand and the Binding Buffer used for bead storage was removed. The beads were resuspended with the master mix and incubated at room temperature with rotation for 30 minutes. After bead binding, the sample was briefly spun down in a mini centrifuge and placed on a magnet stand for 1 minute. The supernatant was removed, and the sample was briefly spun down a second time and returned to the magnet stand for removal of residual supernatant. The sample was then resuspended in 100 μl/sample volume of 1X NEBuffer 2 (NEB) supplemented with Triton X-100 to 0.1% and transferred to a new tube. If the master mix still contained multiple sample volumes at this point, it was divided into individual 100 μl samples. Samples were incubated at room temperature with rotation for 5 minutes, briefly spun down, and placed on a magnet stand for 1 minute. The supernatant was removed, and the samples were briefly spun down a second time and returned to the magnet stand to remove residual supernatant. The beads were then resuspended in 100 μl of Extension Master Mix, which varied by experiment.

100 μl extension reactions were performed with Klenow Fragment (3’→5’ exo-) (NEB) (Figures 2B, 3C), Klenow Fragment (NEB) (Figures 3C, 3D, 4C), or T4 DNA Polymerase (NEB) (Figure 3C). Klenow Fragment (3’→5’ exo-) reactions contained 1X NEBuffer 2, 0.2 mM dNTPs, and 0.1 U/μl Klenow Fragment (3’→5’ exo-) and were incubated at 25 °C for 15 minutes followed by 37 °C for 15 minutes. Klenow Fragment reactions contained 1X NEBuffer 2, 0.2 mM dNTPs, and 0.05 U/μl Klenow Fragment, and were incubated at 25 °C for 20 minutes. T4 DNA Polymerase reactions contained 1X NEBuffer 2.1 (NEB), 0.2 mM dNTPs and 0.03 U/μl T4 DNA Polymerase and were incubated at 12 °C for 20 minutes. All reactions were performed in 200 μl thin-walled tubes on a thermal cycler block. Following extension, the reactions were transferred to a pre-chilled 1.7 ml microcentrifuge tube, placed on a magnet stand, and the supernatant was removed.

**Figure 1.**
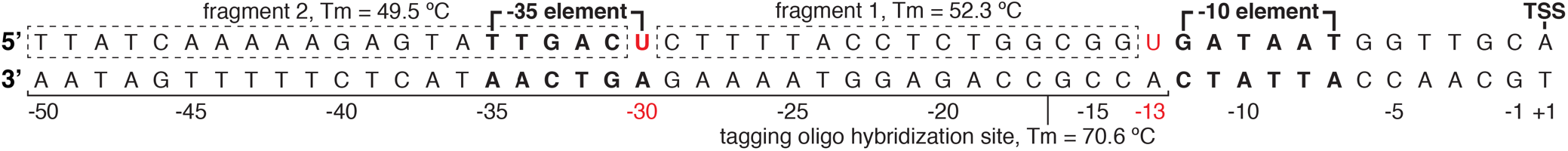
Overview of the deoxyuridine-modified P_RA1_ σ70 promoter. The sequence of the P_RA1_ σ^70^ promoter is annotated to indicate the transcription start site (TSS), −10 element, −35 element, the location of dU nucleotides, and the fragments and tagging oligo hybridization site that result from dU excision. The Tm of fragments 1 and 2 were calculated using the Integrated DNA Technologies OligoAnalyzer Tool with the settings: Oligo conc. = 0.005μM, Na^+^ conc.= 0 mM, Mg^++^ conc. = 10 mM, dNTPs conc. = 1 mM. The Tm of the tagging oligo hybridization site was calculated using the settings: Oligo conc. = 0.05μM, Na^+^ conc.= 0 mM, Mg^++^ conc. = 10 mM, dNTPs conc. = 1 mM. These settings approximate the conditions of the experiment.

**Figure 2.**
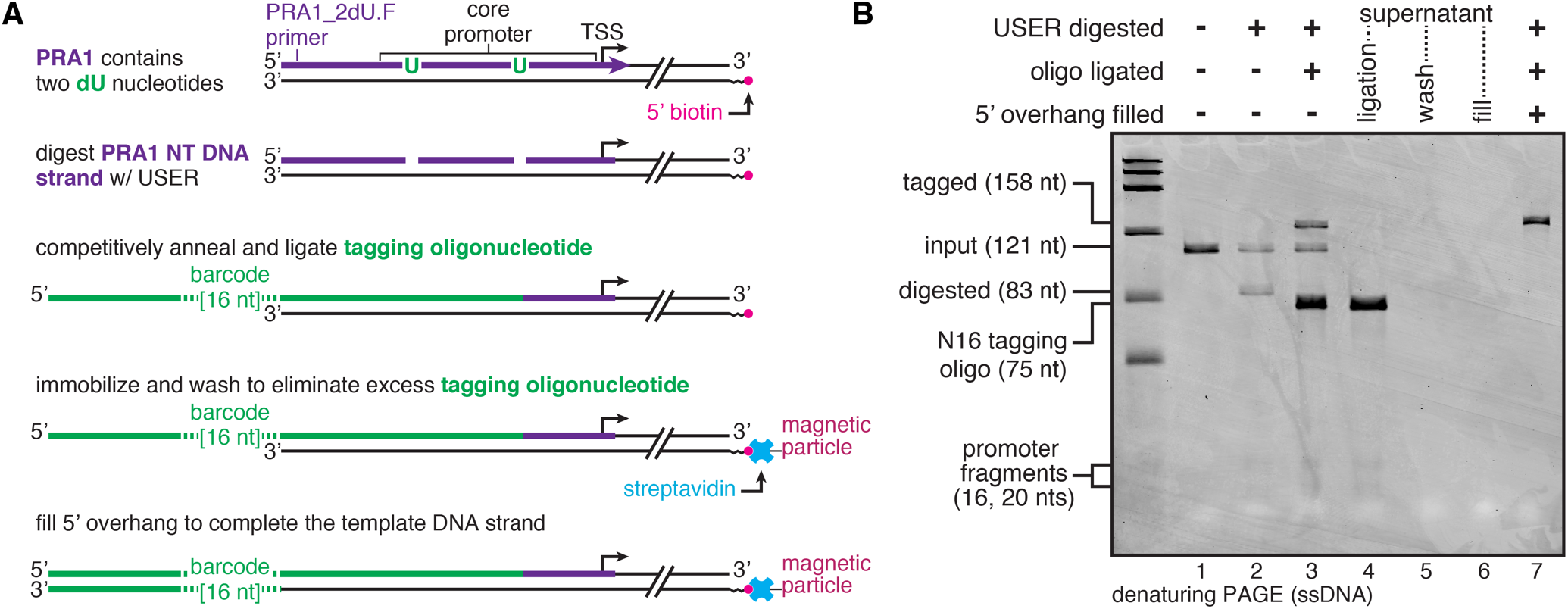
dUX-tagging of blunt dsDNA with streptavidin-coated magnetic bead clean-up. **(A)** Overview of the dUX-tagging procedure when streptavidin-coated magnetic beads are used to deplete excess tagging oligo. **(B)** Denaturing PAGE analysis of dUX-tagging intermediate fractions that were collected from a single pooled reaction when performing the procedure shown in (A). The denaturing PAGE gel used the Low Range ssRNA Ladder (NEB; Markers: 50, 80, 150, 300, 500, 1000 nts). The linear dsDNA used for this experiment contained a 5’ biotin modification. NT, non-transcribed.

**Figure 3.**
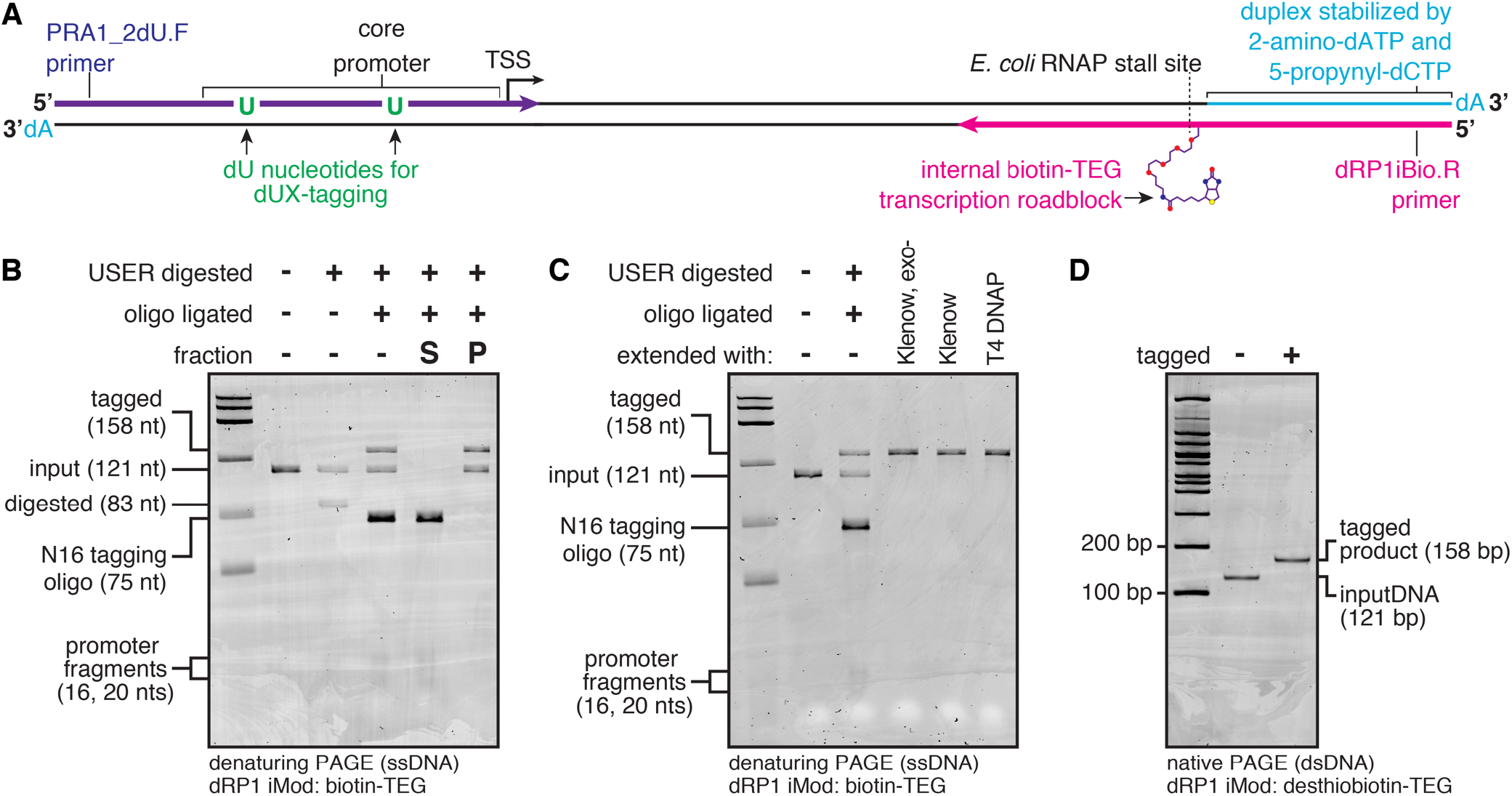
dUX-tagging of internally biotinylated dsDNA with streptavidin-coated magnetic bead clean-up. **(A)** Overview of the internally biotinylated linear dsDNA used to perform the experiments in panels B-D. **(B)** Denaturing PAGE analysis of dUX-tagging intermediate fractions that were collected from a single pooled reaction when performing the procedure shown in Figure 2A up to the excess tagging oligo depletion step. S, supernatant; P, pellet. **(C)** Denaturing PAGE analysis to assess primer extension of tagged internally biotinylated dsDNA using various DNA polymerases. **(D)** Native PAGE analysis of internally desthiobiotinylated DNA that was tagged using the dUX-tagging procedure shown in Figure 2A. Tagged DNA was eluted from streptavidin coated beads without denaturation by supplying excess free biotin to displace desthiobiotin from streptavidin. The denaturing PAGE gels used the Low Range ssRNA Ladder (NEB; Markers: 50, 80, 150, 300, 500, 1000 nts).

**Figure 4.**
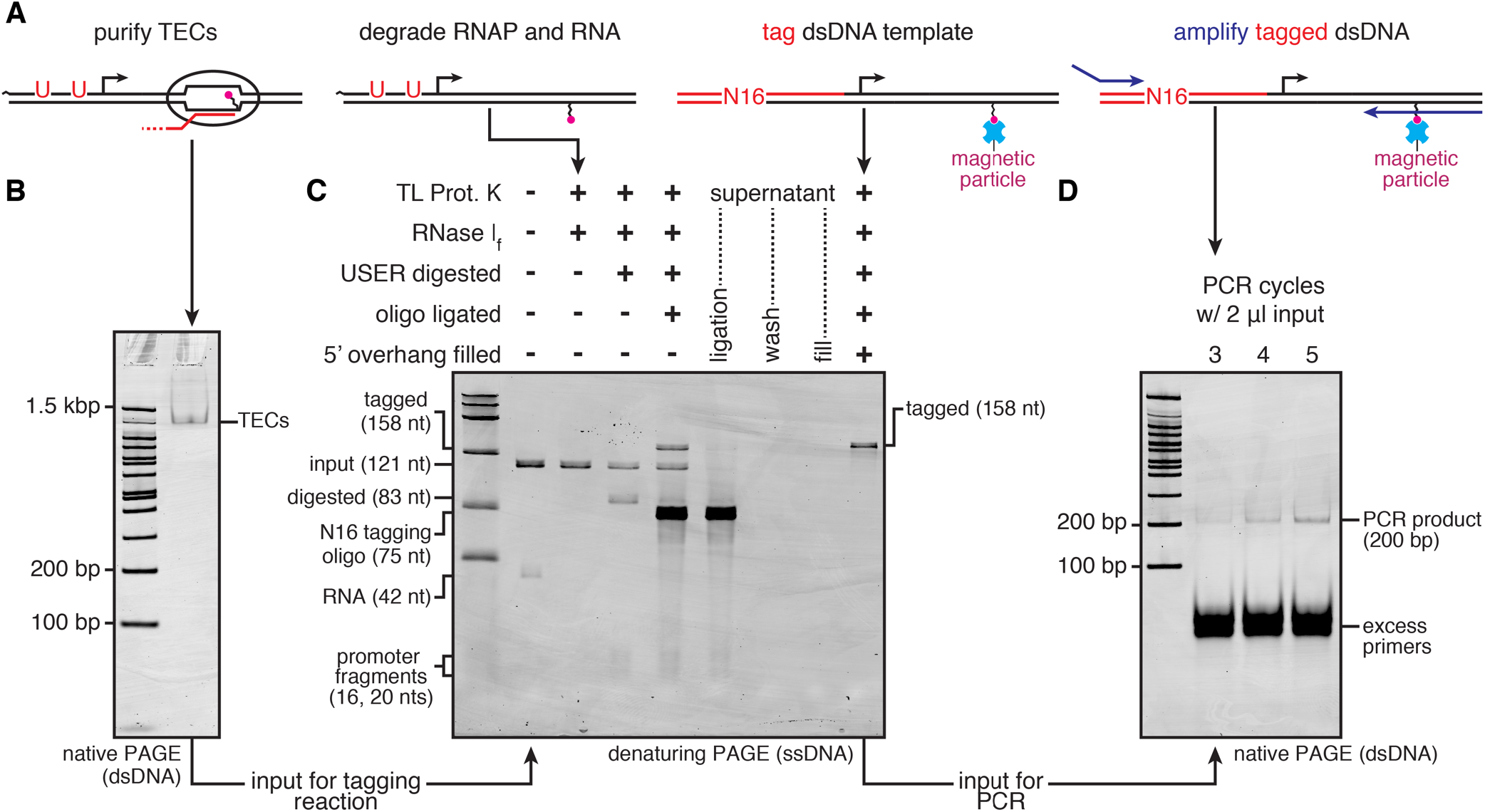
dUX-tagging of DNA recovered from purified TECs. **(A)** Overview of the steps for tagging DNA from purified transcription elongation complexes (TECs). **(B)** TECs were purified using selective photoelution (16) and assessed by native PAGE. **(C)** Purified TECs from (A) were treated with thermolabile proteinase K and RNase I_f_ before performing the dUX-tagging procedure shown in Figure 2A; an intermediate fraction of the combined reaction was collected after each step and assessed by denaturing PAGE. **(D)** 2 μl of 25 μl of tagged DNA from (B) was used as the template for a limited-cycle PCR, and 5 μl of the 25 μl PCR was loaded on the gel. The expected PCR product was visible by native PAGE within 3 amplification cycles. A version of this gel with the grayscale darkened to show trace impurities is presented in Supplementary Figure S1A. The denaturing PAGE gel used the Low Range ssRNA Ladder (NEB; Markers: 50, 80, 150, 300, 500, 1000 nts). NT, non-transcribed; TL Prot. K, Thermolabile Proteinase K.

Immobilized DNA was eluted from streptavidin beads using either a denaturing or non-denaturing procedure: DNA containing either a 5’ or internal biotin modification was recovered by resuspending the bead pellet in 25 μl of 95% deionized formamide and 10 mM EDTA, heating at 100 °C for five minutes, placing the sample on a magnet stand, and collecting the supernatant. DNA containing an internal desthiobiotin was recovered by resuspending the bead pellet in 150 μl of Biotin Elution Buffer (0.5 M Tris-HCl (pH 8.0), 10 mM EDTA (pH 8.0), 10% DMSO, 2 mM D-Biotin), incubating at room temperature with rotation for 15 minutes, placing the sample on a magnet stand, and collecting the supernatant.

#### High-specificity primer extension and tagging oligonucleotide degradation

After the ligation step was completed, 200 ng/sample volume (0.4 μl) of ET SSB (NEB) was added, and T4 DNA ligase was inactivated by incubation at 65 °C for 10 minutes. In the final protocol, each 100 μl primer extension reaction contained 26.9 μl (one sample volume) of the dUX-tagging reaction, 1X ThermoPol Buffer (NEB), 0.2 mM dNTPs, 2.5% deionized formamide, and 0.02 U/μl Vent (exo-) DNA polymerase (NEB). Primer extension was also performed without ET SSB, without formamide, with 0.04 U/μl Vent (exo-) DNA polymerase, with 0.02 U/μl Vent DNA polymerase (NEB) instead of Vent (exo-), and with both Vent (exo-) and 0.02 U/μl *Sulfolobus* DNA polymerase IV (NEB) as indicated (Figures 5B, 6A and 6B). Reactions were aliquoted into 200 μl thin-walled tubes and kept in an aluminum block on ice. The thermal cycler block was pre-heated to 72 °C with a heated lid set to 105 °C. Reactions were transferred to the 72 °C thermal cycler block and incubated for 5 min (or 10/15 min in Figure 6B). The reactions were cooled rapidly by transferring the tubes from the 72 °C thermal cycler block back to the aluminum block on ice.

**Figure 5.**
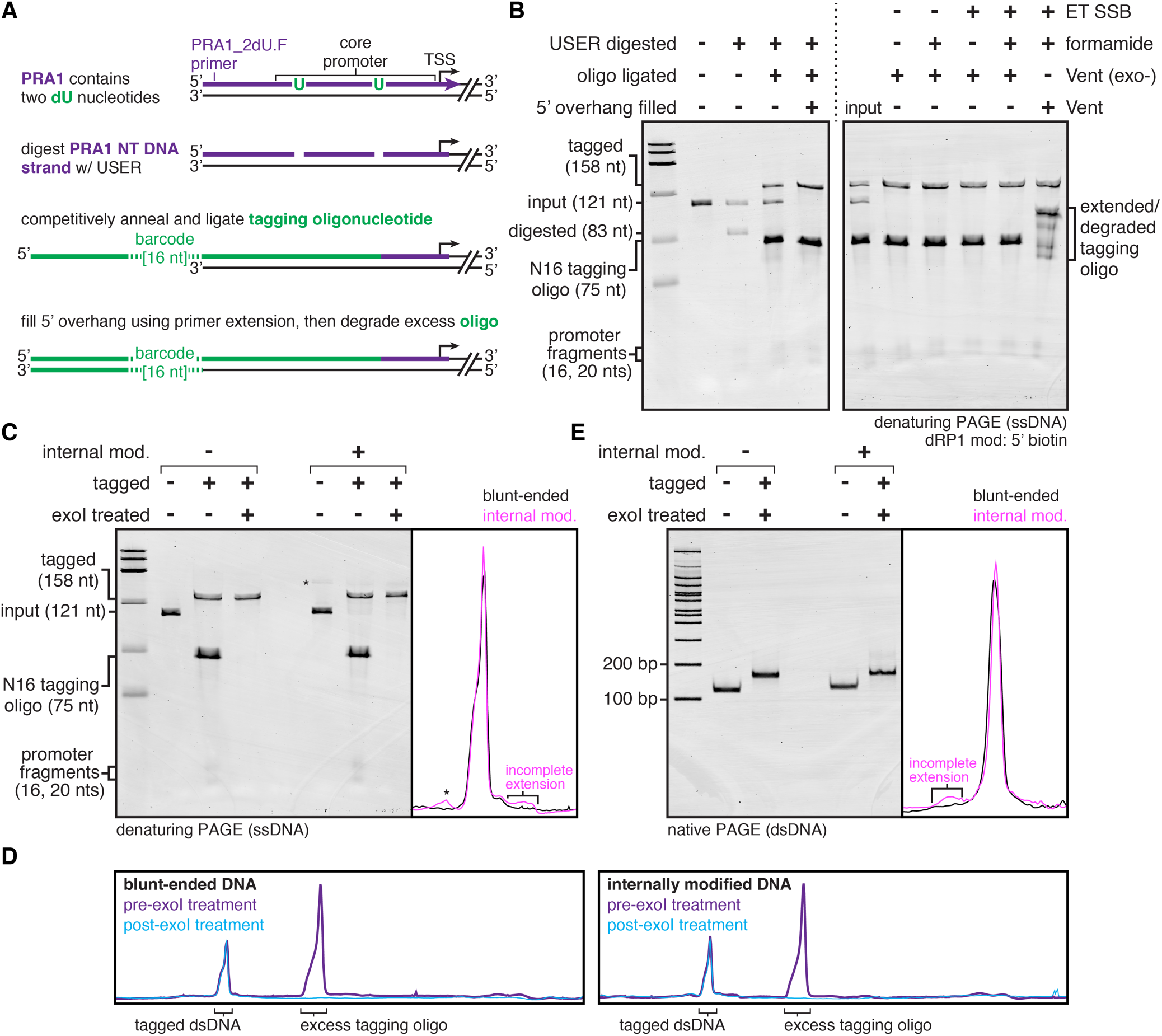
dUX-tagging of dsDNA with exonuclease I clean-up. **(A)** Overview of the dUX-tagging procedure when thermolabile exonuclease I is used to degrade excess tagging oligo. In contrast to the procedure shown in Figure 2A, which was a one-pot reaction through tagging oligo ligation, this version of the dUX-tagging method remains one-pot through the primer extension step. **(B)** Denaturing PAGE analysis of dUX-tagging intermediate fractions that were collected from a single pooled reaction when performing the procedure shown in (A) with blunt-ended DNA. Several primer extension conditions are shown. **(C)** Denaturing PAGE analysis of tagging oligo degradation after the procedure in (A) was applied to blunt and internally biotinylated dsDNA. Incomplete primer extension that occurs with the internally biotinylated DNA is shown in the intensity trace. The asterisk indicates a minor dsDNA product that was present in the internally modified linear dsDNA preparation. **(D)** Intensity traces comparing the pre- and post-exoI treatment samples from (C) for both blunt and internally biotinylated DNA. **(E)** Native PAGE analysis of blunt and internally biotinylated DNA before and after performing the procedure in (A). Incomplete primer extension that occurs with the internally biotinylated DNA is shown in the intensity trace. dsDNA with incomplete primer extension migrates slower than fully double stranded DNA (Supplementary Figure S2). The denaturing PAGE gels used the Low Range ssRNA Ladder (NEB; Markers: 50, 80, 150, 300, 500, 1000 nts). ET SSB, extreme thermostable single-stranded DNA binding protein; exoI, thermolabile exonuclease I.

**Figure 6.**
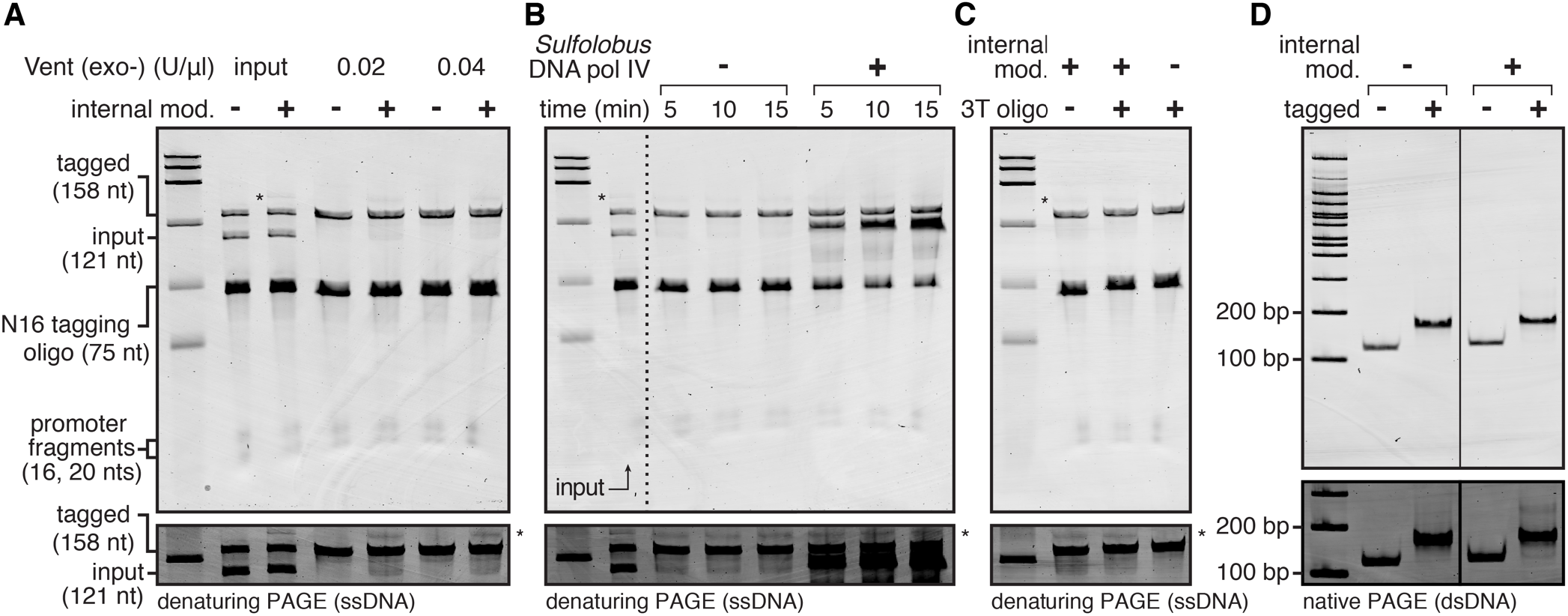
Primer extension of tagged DNA using variable reaction conditions. **(A)** Denaturing PAGE analysis of primer extension reactions using tagged blunt and internally modified DNA and a variable amount of Vent (exo-) DNA polymerase. **(B)** Denaturing PAGE analysis of primer extension reactions using tagged, internally modified DNA with or without *Sulfolobus* DNA polymerase IV for variable reaction times. **(C)** Denaturing PAGE analysis of blunt and internally modified DNA that was tagged with either the VRA5_16N_PRA1m12 or VRA5_16N_3T_PRA1m12 (3T oligo) tagging oligos (Supplementary Table S1). **(D)** Native PAGE analysis of blunt and internally modified DNA that was tagged with the 3T oligo from (C). In each panel, a cut-out of the substrate DNA band is shown with a darkened grayscale setting to better visualize incomplete primer extension products. In panels A, B, and C, the asterisks indicate a minor dsDNA product that was present in the internally modified linear dsDNA preparation. The denaturing PAGE gels used the Low Range ssRNA Ladder (NEB; Markers: 50, 80, 150, 300, 500, 1000 nts).

Before exonuclease treatment, PCRs were purified by phenol:chloroform extraction and ethanol precipitation to remove protein and formamide. An equal volume (100 μl) of Tris (pH 8) buffered phenol:chloroform:isoamyl alcohol (25:24:1, v/v) was added to each sample, the sample was mixed by vortexing and inversion and centrifuged at 18,500 x g and 4 °C for 5 min. The aqueous phase was collected and transferred to a new tube. DNA was precipitated by adding 0.1 volumes (10 μl) of 3 M NaOAc (pH 5.5), 3 volumes (300 μl) of 100% ethanol, and 1.5 μl of GlycoBlue Coprecipitant to each sample. Samples were chilled at −70 °C for 30 min and centrifuged at 18,500 x g and 4 °C for 30 min. After removing the supernatant, the samples were washed once by adding 1 ml of cold 70% ethanol, inverting the tube several times, and centrifuging for 5 min. The supernatant was removed and the samples were centrifuged again briefly, and residual supernatant was removed. Pelleted DNA was resuspended in 30 μl of NEBuffer 3.1 (NEB) and 0.5 μl of Thermolabile Exonuclease I (NEB) was added. Samples were transferred to 200 μl thin-walled tubes and incubated on a thermal cycler block set to 37 °C with a heated lid set to 45 °C for 4 minutes. Samples were placed in an aluminum block on ice and the thermal cycler was set to 80 °C with the heated lid set to 105 °C. Samples were transferred to the hot block and incubated at 80 °C for 1 minute to heat-inactivate Thermolabile Exonuclease I.

#### Collection and processing of intermediate dUX-tagging fractions

Intermediate dUX-tagging fractions were collected as follows: Unprocessed control fractions were collected by transferring 24.5 μl of the initial master mix to 125 μl of Stop Solution (0.6 M Tris-HCl (pH 8.0), 12 mM EDTA (pH 8.0)) before Thermolabile USER^®^ II Enzyme was added. USER digested fractions were collected by transferring 25 μl of master mix to 125 μl of Stop Solution after the dU excision reaction. Oligo ligated fractions were collected by transferring 26.5 μl of master mix to 125 μl of Stop Solution after the tagging oligonucleotide ligation. For dUX-tagging with streptavidin-bead-based clean-up, wash fractions were collected by transferring a wash supernatant volume that corresponded to one sample to Stop Solution for a total volume of 150 μl, fully tagged fractions that were eluted from beads by heat denaturation were collected by transferring the 25 μl supernatant to 125 μl of Stop Solution, and fully tagged fractions that were eluted in 150 μl of Biotin Elution Buffer were collected directly. For dUX-tagging with exonuclease I clean-up, 100 μl primer extension reactions were collected by mixing with 50 μl of Stop Solution supplemented with 500 mM EDTA (pH 8.0) so that the final EDTA concentration was 10 mM, and thermolabile exonuclease I reactions were collected by mixing with 125 μl of Stop Solution.

All fractions were processed by phenol:chloroform extraction and ethanol precipitation as described above in the section *High-specificity primer extension and tagging oligonucleotide degradation* with the volume of each reagent adjusted for a 150 μl sample volume. In Figures 2 and 3, samples for denaturing PAGE were resuspended in 32 μl of Formamide Loading Dye+XC (90% (v/v) deionized formamide, 1X Transcription Buffer (20 mM Tris-HCl (pH 8.0), 50 mM KCl, 1 mM DTT, 0.1 mM EDTA (pH 8.0)), 0.025% (w/v) bromophenol blue, 0.025% (w/v) xylene cyanole FF) and half the sample was used for denaturing PAGE. In Figures 4, 5, and 6, samples for denaturing PAGE were resuspended in 16 μl of Formamide Loading Dye (90% (v/v) deionized formamide, 1X Transcription Buffer, 0.025% (w/v) bromophenol blue) and the entire sample was used for denaturing PAGE. Samples for native PAGE were resuspended in 10 μl of 10 mM Tris-HCl (pH 8.0) and 2 μl of 6X DNA loading dye (30% (v/v) glycerol, 0.025% (w/v) bromophenol blue).

### dUX-tagging with DNA from purified TECs

TECs were prepared in bulk exactly as described previously (16). A 15 μl aliquot was removed and assessed by EMSA exactly as described previously (16) and a 25 μl ‘Input’ fraction was mixed with 125 μl of Stop Solution, phenol:chloform extracted and set up for ethanol precipitation as described above in the section *Collection and processing of intermediate dUX-tagging fractions*. All subsequent incubations were performed in a thermal cycler. 150 μl of the remaining sample was mixed with 6 μl (1 μl per 25 μl sample volume) of Thermolabile Proteinase K (NEB), incubated at 37 °C for 30 minutes, and then at 65 °C for 20 minutes. 3 μl (0.5 μl per sample volume) of RNase I_f_ (NEB) was added and the sample was incubated at 37 °C for 15 minutes, and then at 70 °C for 20 minutes. Together these steps degraded RNAP and RNA, and the sample was stored at −20 °C overnight. The next day, 26.5 μl was removed and mixed with 125 μl of Stop Solution. The remaining sample was mixed with 12.5 μl (2.5 μl per sample volume) of 10X T4 DNA ligase Buffer and 2.5 μl (0.5 μl per sample volume) of Thermolablie USER^®^ II Enzyme and processed exactly as described above in the sections *USER digestion, tagging oligonucleotide annealing, and DNA ligation* and *Bead-based tagging oligonucleotide clean-up and low-temperature primer extension* through the Klenow Fragment primer extension step. After each enzymatic processing step a reaction fraction (with volume adjusted for added components) was collected as described above in the section *Collection and processing of intermediate dUX-tagging fractions*. All intermediate fractions were phenol:chloroform extracted and ethanol precipitated as described above, and the pellet was resuspended in 16 μl of Formamide Loading Dye for denaturing PAGE. After all intermediate fractions were collected, this procedure yielded two reaction volumes of bead-immobilized tagged DNA. The beads were washed once with 1 ml of 10 mM Tris-HCl (pH 8.0) supplemented with 0.01% Triton X-100, and resuspended in 25 μl per sample volume of this same buffer for storage at −20 °C.

PCR of the tagged DNA was performed in 25 μl reactions in 200 μl thin-walled PCR tubes containing 1X Q5^®^ Buffer (NEB), 1X Q5^®^ GC Enhancer (NEB), 0.2 mM dNTPs, 0.25 μM primer RPI1 (Supplementary Table S1), 0.25 μM primer dRP1_NoMod.R (Supplementary Table S1), 2 μl bead-immobilized tagged DNA, and 0.02 U/μl Q5^®^ High-Fidelity DNA Polymerase (NEB). Reactions were amplified using the thermal cycling program: 98 °C for 30 s, [98 °C for 10 s, 69 °C for 20 s, 72 °C for 20 s] x N cycles (where N = 3, 4, or 5 cycles as indicated in Figure 4D), hold at 12 °C. After amplification, reactions were transferred to 1.7 ml microcentrifuge tubes and placed on a magnet stand to pellet the beads. The supernatant was collected, and 5 μl of supernatant was mixed with 1 μl of 6X BPB Only SDS DNA Loading Dye (30% (v/v) glycerol, 10 mM Tris-HCl (pH 8.0), 0.48% (w/v) SDS, 0.01% (w/v) bromophenol blue) for electrophoresis by native PAGE.

### Gel electrophoresis

Denaturing urea-PAGE, native PAGE, and transcription elongation complex EMSAs were performed exactly as described previously (16). Denaturing urea-PAGE gels used the Low Range ssRNA Ladder (NEB). The resulting gels were stained with SYBR Gold and scanned on a Sapphire™ Biomolecular Imager (Azure Biosystems) using the 488 nm/518BP22 setting exactly as described previously (16).

### Quantification

Quantification of incomplete primer extension was performed using ImageJ 1.51s by plotting each lane, drawing a line at the base of the full-length product band and incomplete product smear, and determining the area of the closed sections. Primer extension efficiency was calculated by dividing the intensity of the complete product band by the sum of the intensities of the complete product band and incomplete product smear.

## Results

### Overview of the dUX-tagging procedure

The goal of dUX-tagging is to append sequence to a linear dsDNA molecule by swapping a user-specified 5’-end segment of one DNA strand with a synthetic oligonucleotide. The sections below describe two implementations of this procedure that share four common steps: 1) Excise dU nucleotides from the substrate dsDNA to generate a 3’ overhang. 2) Competitively anneal and ligate a tagging oligo to the substrate DNA. For all experiments in this work the tagging oligo contained an N16 randomized segment flanked by constant DNA sequences to ensure that dUX-tagging is compatible with molecular barcoding applications (Supplementary Table S1). 3) Remove excess tagging oligo. 4) Fill the 5’ overhang that results from tagging oligo ligation using primer extension. This approach is sequence-independent because the 3’ overhang used to anneal the tagging oligo is defined by the location of dU nucleotides rather than a DNA sequence motif. Both implementations of dUX-tagging are one-pot reactions through the tagging oligo ligation step to minimize variability from sample handling. The primary difference between the two implementations is how excess tagging oligo is removed from the reaction. In the first procedure, the substrate DNA is biotinylated and can be immobilized so that excess tagging oligo is washed away before primer extension. In the second procedure, primer extension is performed in the presence of the tagging oligo, which can then be degraded by exonuclease I treatment.

To illustrate how dUX-tagging functionality can be integrated with a biologically active DNA element, I developed the procedures below using a custom phage-derived σ^70^ promoter called P_RA1_ (16) as the tagging site. dU substitutions were previously added to P_RA1_ at positions −13 (one nucleotide upstream of the −10 element hexamer) and at −30 (the sixth nucleotide in the −35 element hexamer) to facilitate a promoter-specific ‘USER enzyme footprinting assay’ (Figure 1) (16). Crystallographic structures of bacterial promoter complexes suggested that dT→dU base changes at these positions should not interfere with function (19–22), and I previously showed that the dU-modified P_RA1_ promoter can be saturated by open promoter complexes and used for *in vitro* transcription (16). In addition to maintaining promoter function, these positions were selected so that to the Tm of the DNA fragments that form upon deoxyuridine excision is approximately 20 °C lower than the Tm the entire 3’ overhang (Figure 1). A complete description of the design considerations that were taken into account when embedding a dUX-tagging site in the P_RA1_ promoter is presented in the Discussion.

### Strategy 1: dUX-tagging with streptavidin-coated magnetic bead clean up

When the dsDNA substrate for dUX-tagging can contain a biotin modification, excess tagging oligo can be removed by performing a straightforward immobilization and washing procedure after the tagging reaction, as follows (Figure 2A): 1) dU nucleotides are excised by digestion with the thermolabile USER II enzyme mixture (Figure 2B, lane 2). 2) The tagging oligo is competitively annealed to the resulting 3’ overhang by heating to 70 °C and slowly cooling to 25 °C; thermolabile USER II enzyme is inactivated at this step. 3) The tagging oligo is ligated to the substrate DNA by T4 DNA ligase (Figure 2B, lane 3). T4 DNA ligase is then heat inactivated. After this step, the DNA has been tagged on one strand and contains a 5’ overhang (Figure 2A). 4) The DNA is immobilized on streptavidin-coated magnetic beads, the binding supernatant is removed, and the beads are washed once. All tagging oligo is removed by this procedure and the DNA substrate is retained on the beads (Figure 2B, lanes 4 and 5). 5) The 5’ overhang is filled by primer extension with Klenow Fragment (3’→5’ exo-) (Figure 2B, lane 7). Removing the supernatant after primer extension functions as a third wash (Figure 2B, lane 6). The beads can then be washed and resuspended with 10 mM Tris-HCl (pH 8.0) and stored at −20 °C. Each step of this procedure is efficient, and no reaction side products were observed.

### Application of dUX-tagging to internally biotinylated DNA

The procedure above was developed using blunt-ended DNA that did not contain any internal modifications other than the dU nucleotides required for dUX-tagging. To show that dUX-tagging is compatible with a more complex DNA substrate, I barcoded a DNA template that contains an internal biotin-TEG modification, which can be used as an *E*. *coli* RNA polymerase (RNAP) transcription stall site (16,23) (Figure 3A). DNA that contains an internal biotin-TEG modification is prepared using a procedure that requires translesion DNA synthesis by *Sulfolobus* DNA polymerase IV (24), which has two consequences: First, *Sulfolobus* DNA polymerase IV bypasses the internal biotin-TEG modification without incorporating a nucleotide (23). This distorts the DNA duplex and may reduce the thermal stability of the DNA helix downstream of the modification site, which typically contains ~30 base pairs. This was addressed during linear dsDNA preparation by performing translesion synthesis using a dNTP mixture in which dATP and dCTP were substituted with the thermostability-enhancing dNTPs 2-amino-dATP and 5-propynyl-dCTP (Figure 3A). Second, *Sulfolobus* DNA polymerase IV leaves a 1-2 nt dA (2-amino-dATP, in this case) overhang on the DNA 3’ ends (25), which may interfere with primer extension (Figure 3A). Control experiments showed that DNA containing an internal biotin-TEG modification is efficiently tagged and separated from excess tagging oligo using streptavidin beads (Figure 3B). Primer extension to fill the 5’ overhang that results from tagging oligo ligation went to completion with Klenow Fragment (3’→5’ exo-), Klenow Fragment, and T4 DNA polymerase (Figure 3C). Native PAGE analysis of a dUX-tagging reaction that was performed with an internally desthiobiotinylated substrate DNA showed that tagging is efficient and does not result in any detectable side products (Figure 3D).

### Application of dUX-tagging to DNA derived from transcription elongation complexes

In the experiments above, dUX-tagging was applied to pure DNA. To show that dUX-tagging can be applied to a more complex sample, I barcoded DNA derived from purified *E. coli* RNAP transcription elongation complexes (TECs) (Figure 4A). TECs were positioned at an internal biotin-TEG stall site and purified by selective photoelution from magnetic beads (16) (Figure 4A, B). The purified TECs were pre-processed for dUX-tagging by treating with Thermolabile Proteinase K to degrade RNAP, heat-inactivating Thermolabile Proteinase K, degrading the RNA transcript by treating with RNase I_f_, and heat-inactivating RNase I_f_ (Figure 4A, C). The resulting DNA was barcoded efficiently using the dUX-tagging procedure shown in Figure 2A (Figure 4A, C). Amplification using the standard Illumina RNA PCR Index Primer (which anneals to constant sequence appended by the tagging oligonucleotide) and an extended version of the Illumina RNA PCR Primer (which anneals to a constant region in the substrate DNA) yielded the expected PCR product (Figure 4A, D and Supplementary Table S1). Trace amounts of discrete slower and faster migrating products were detectable (Supplementary Figure 1A). However, these non-target products were not amplifiable and therefore mostly likely correspond to DNA duplexes that contain one strand from the original tagged DNA sample (Supplementary Figure 1B). Together, these analyses show that the dUX-tagging procedure is compatible with DNA that was recovered from a complex sample that contained protein and RNA components. In this case, no additional DNA purification was required for efficient DNA tagging.

### Strategy 2: dUX-tagging with exonuclease I clean up

In some cases the dsDNA substrate for dUX-tagging will not contain a biotin modification so that excess tagging oligonucleotide cannot be removed using an immobilization-based approach. To address this, I implemented a second dUX-tagging procedure in which primer extension is performed in the presence of the tagging oligo, which is later degraded by exonuclease I (Figure 5A, B). In this procedure, dU excision, tagging oligo annealing, and ligation are performed exactly as described above. When excess tagging oligo is depleted immediately after ligation, primer extension can be performed at low temperature because primer dimer formation is not a concern (Figures 2B, 3C, 4C). When performing primer extension in the presence of the tagging oligo, primer dimer formation is a substantial concern due to the N16 barcode region. To address this, primer extension was performed as follows: First, a commercially-available thermostable single-stranded DNA binding protein (ET SSB, New England Biolabs) was added to the sample before heat-inactivating T4 DNA ligase to reduce primer dimer formation (26). Second, the primer extension reaction contained 2.5% formamide, which can improve oligonucleotide hybridization specificity (27). Third, Vent (exo-) DNA polymerase was used so that primer extension is performed at 72 °C (28). The 3’→5’ exonuclease activity of Vent DNA polymerase caused primer dimer to form even in the presence of specificity-enhancing additives (Figure 5B). Fourth, heating and cooling was performed rapidly by moving samples from a chilled aluminum block to a pre-heated thermal cycler block and immediately returning the samples to the chilled aluminum block after the five minute reaction was complete. The purpose of this manipulation was to limit the possibility of primer dimer formation at intermediate temperatures during sample heating and cooling. In my laboratory, including ET SSB and 2.5% formamide in the primer extension reaction was not necessary to avoid primer dimer formation (Figure 5B). However, during preliminary development of this procedure in a separate laboratory setting these additives were required to avoid primer dimer. The use of ET SSB and formamide is therefore not strictly required, however they are not detrimental to the primer extension reaction and likely improve the robustness of the procedure to variations in instrumentation and sample handling. After primer extension, the tagged DNA was phenol:chloroform extracted and ethanol precipitated to remove protein and formamide. Excess tagging oligo was degraded by treatment with thermolabile exonuclease I, which was subsequently heat-inactivated (Figure 5C, E). No meaningful loss of DNA resulted from these steps (Figure 5C, D).

When dUX-tagging was applied to a dsDNA substrate that contained an internal biotin-TEG modification ~5% of the DNA did not undergo primer extension (Figure 5C, E). Several attempts to improve primer extension yield by adjusting reaction conditions were unsuccessful: Increasing the concentration of Vent (exo-) DNA polymerase or the duration of primer extension had no effect (Figure 6A, B). This suggests that the unextended dsDNA is not a substrate for extension by Vent (exo-) DNA polymerase. Importantly, increasing Vent (exo-) concentration or the reaction time did not cause the formation of primer dimer (Figure 6A, B). Including *Sulfolobus* DNA polymerase IV caused the time-dependent formation of substantial amounts of primer dimer (Figure 6B). Using a tagging oligo that contained three dT nucleotides to account for the inferred 2-amino-dA 3’ overhang (Figure 3A) did not improve primer extension efficiency (Figure 6C, D and Supplementary Table S1). Nonetheless, because the dUX-tagging procedure is effectively quantitative through the tagging oligo ligation step, essentially every dsDNA molecule is tagged on at least one strand so that the tagging efficiency for internally modified DNA was >97% when considered on a per-strand basis.

## Discussion

In this work I have developed two variations of a site-specific and sequence-independent method for linear dsDNA tagging, called dUX-tagging. This approach uses dU excision to generate a 3’ overhang so that a complementary oligonucleotide can be annealed, ligated, and used as a template to extend the substrate dsDNA molecule. In this way, arbitrary sequences can be appended to linear dsDNA in a concise and efficient procedure that requires only commercially available reagents.

The utility of dUX-tagging is derived from three properties: First, dUX-tagging is sequence-independent because the tagging site is specified by dU bases rather than a sequence motif. This property ensures that tagging is compatible with complex DNA sequence libraries because DNA cutting does not occur at any sites other than the dU nucleotides. Furthermore, tagging sites can be integrated with biologically active DNA elements, such as the P_RA1_ σ^70^ promoter used for method development, by identifying minimally perturbative dT→dU substitutions. Second, both variations of the dUX-tagging procedure are performed as a one-pot reaction at least through the tagging oligo ligation step and can remain one-pot through primer extension if desired (Figure 5). By design, this enables differentially tagged samples to be pooled after the one-pot tagging reaction so that any downstream processing steps are uniform. Third, both variations of dUX-tagging are efficient. When excess tagging oligo was depleted immediately after ligation so that primer extension could be performed at low temperature with proofreading DNA polymerases, tagging was effectively complete for both blunt-ended and internally modified DNA (Figures 2 and 3). When primer extension was performed immediately after ligation using specificity-enhancing reaction conditions, tagging was effectively complete for blunt-ended DNA and >97% efficient on a per-strand basis for internally modified DNA due to incomplete extension of ~5% of the tagged DNA (Figure 5). Because primer extension cannot be performed with a proofreading DNA polymerase when tagging oligo is present in the reaction (Figure 5B), it is crucial to benchmark tagging efficiency for any DNA that is not blunt-ended. Importantly, even when primer extension was incomplete, DNA tagging was effectively complete on a per-DNA-duplex basis due to the high efficiency of the one-pot tagging reaction.

The most crucial consideration when designing a dUX-tagging site is the location of the dU nucleotides. Three criteria should be prioritized: First, dT→dU substitutions should be minimally perturbative to any underlying DNA elements. Second, the placement of dU nucleotides should promote efficient excision. A systematic analysis of the context-dependence of *E*. *coli* uracil-DNA glycosylase activity found that the rate of uracil excision increases when the nucleotide 3’ to dU is dG or dC, and decreases when the nucleotide 3’ to dU is dT (29). Importantly, all sites were eventually processed over the course of a 30 minute reaction (29). In addition, crystal structures of DNA-bound Uracil-DNA glycosylase (30) and endonuclease III (31) suggest that, if possible, dU nucleotides should be flanked by at least 8 bp to avoid steric clashes that would prevent multiple dU sites from being processed simultaneously. Third, the 3’ overhang generated by dU excision must be suitable for annealing a complementary oligonucleotide. When embedding a dUX-tagging site in the P_RA1_ σ^70^ promoter, these three criteria were taken into account in the following ways: −13U was selected because it is the nucleotide immediately upstream of the −10 element hexamer, which is not known to contribute meaningfully to *E*. *coli* σ^70^ promoter function (32), and because it is flanked by GC base pairs. Similarly, −30U was selected despite its position as the sixth nucleotide in the −35 element hexamer because the nucleotide at this position in the non-transcribed DNA strand does not directly contact factor (19) and because it is flanked by GC base pairs (although −29 was changed from A to C specifically for this purpose when P_RA1_ was derived from λP_R_). In addition, −13U and −30U are both flanked by >8 bp and are separated by 16 bp. Finally, the Tm of the 38 nt 3’ overhang was ~20 °C higher than that of the 16 and 20 nt ssDNA fragments that were produced by dU excision in order to promote efficient tagging oligo hybridization. With the guidelines above, incorporation of a dUX-tagging site into the P_RA1_ promoter did not require any optimization.

The dUX-tagging procedures described here have two primary limitations. First, dUX-tagging requires dU nucleotides at defined DNA positions and will therefore be most useful when applied to linear dsDNA that was prepared by PCR for *in vitro* applications. Second, although it was straightforward to embed a dUX-tagging site in the P_RA1_ promoter, some sequences may pose a more substantial challenge. For example, tagging oligonucleotides that are particularly prone to primer dimer formation might be incompatible with even high-specificity primer extension conditions and need to be depleted immediately after ligation. This type of potential pitfall can be avoided by assessing each new dUX-tagging site for efficiency and side product formation using the native and denaturing PAGE assays used to develop the method here.

## Data availability

All data are contained in the manuscript as plotted values or representative gels. Source TIFF files are available from the corresponding author (E.J.S.) upon request.

## Acknowledgements

I am grateful to J.T. Lis (Cornell University) and J.B. Lucks (Northwestern University) for allowing me to conduct preliminary experiments for this work in their laboratories.

## Materials Included

Figure S1. PCR of tagged DNA from TECs with darkened grayscale to show trace impurities
Figure S2. Migration pattern of tagged, internally modified DNA with a 5’ overhang.
Table S1. Oligonucleotides used in this study.
Table S2. Linear dsDNA prepared for this study.
Table S3. Linear dsDNA sequences.

**Figure S1.**
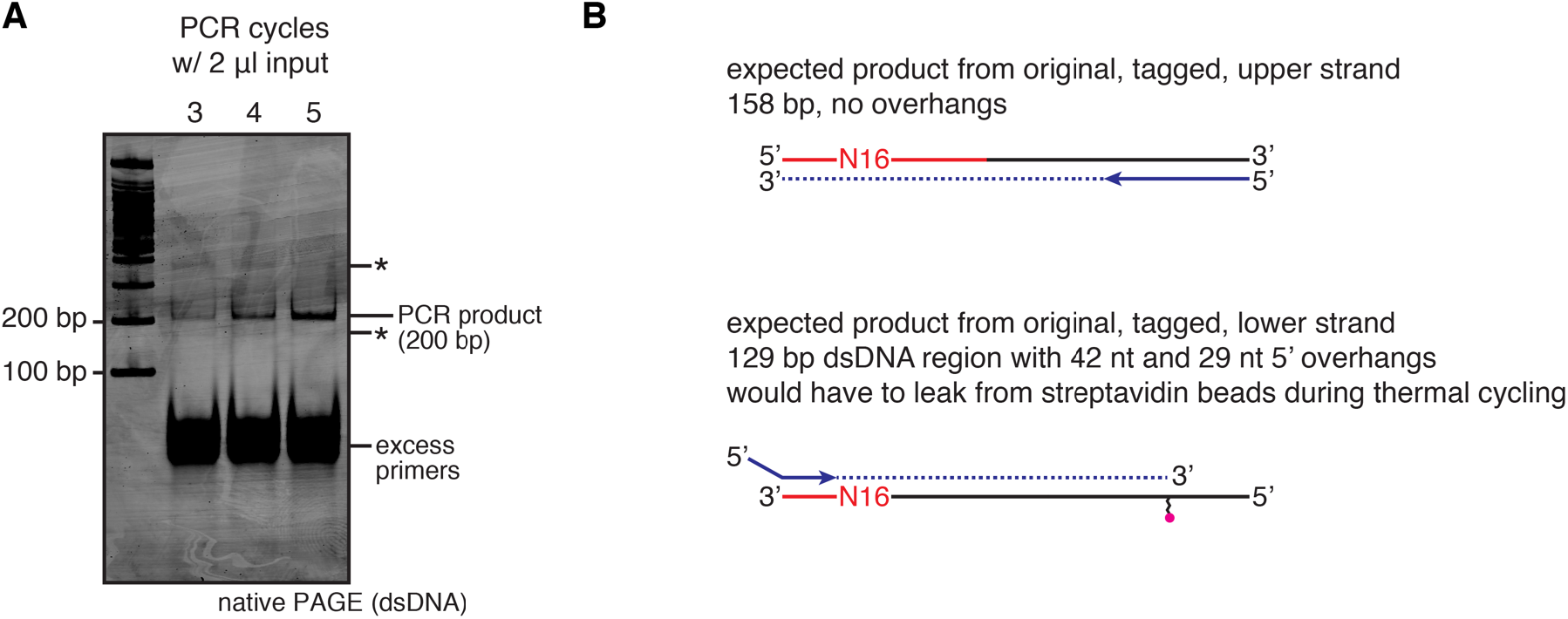
PCR of tagged DNA from TECs with darkened grayscale to show trace impurities. **(A)** The gel from Figure 4D is shown here with the grayscale adjusted to show minor bands that appear during PCR. These products, which are denoted by asterisks, are not amplifiable, whereas the expected PCR product doubles with each amplification cycle. **(B)** Possible identities of the trace impurities from (A).

**Figure S2.**
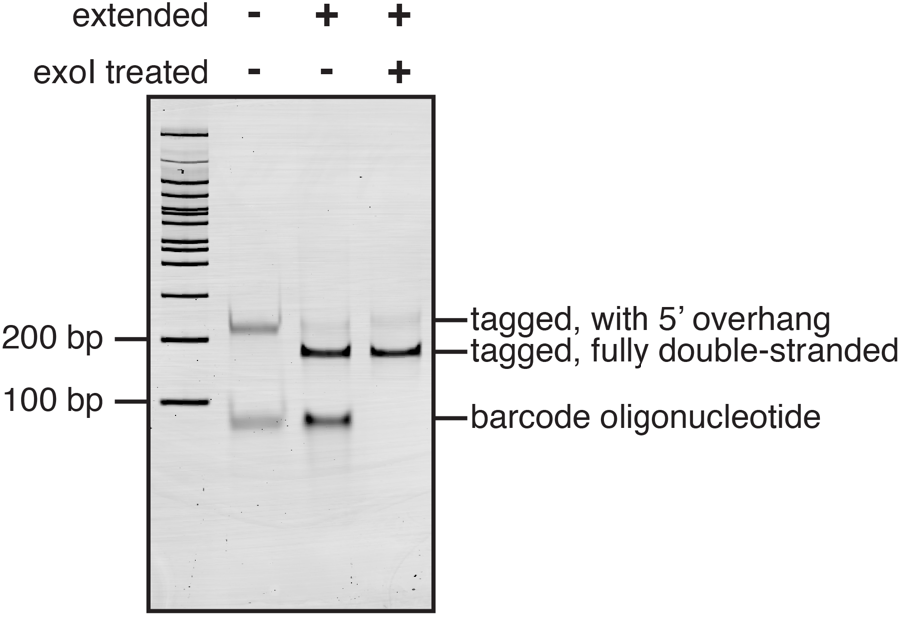
Migration pattern of tagged, internally modified DNA with a 5’ overhang. Native PAGE of tagged, internally modified DNA after ligation, after primer extension, and after exonuclease I treatment.

~~~
/ideSBioTEG/: internal desthiobiotin-triethylene glycol
/iBiotinTEG/: internal biotin-triethylene glycol
/iEth-dA/: internal etheno-dA
/ideoxyU/: internal deoxyuridine
/5PCBio/: 5’ photocleavable biotin
/5bioSG/: “standard” 5’ biotin
~~~

**Table S1.**
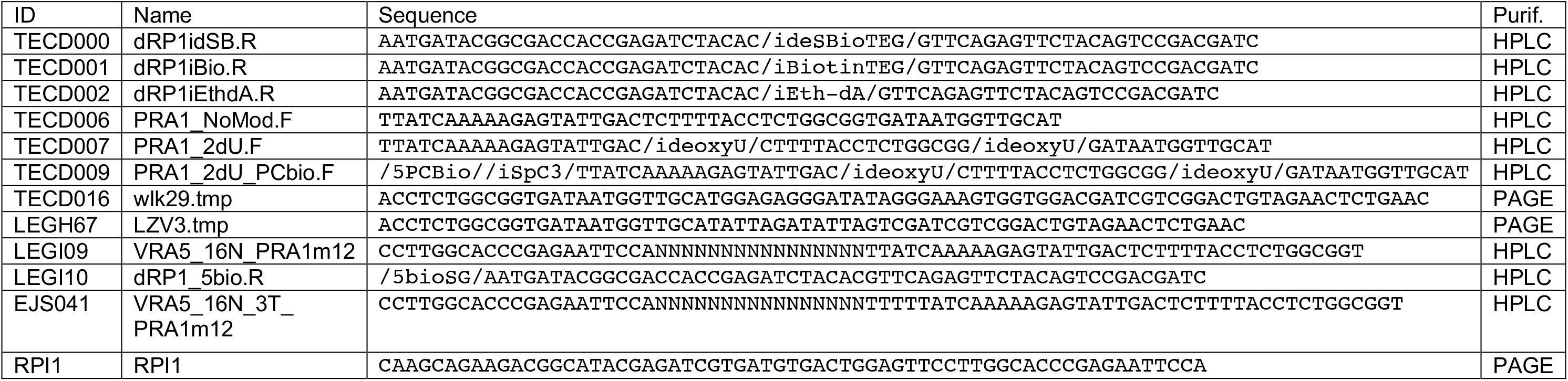
Oligonucleotides used in this study. Below is a table of oligonucleotides used for the preparation of linear dsDNA templates and for the dUX-tagging procedure. The modification codes defined below are used for compatibility with Integrated DNA Technologies ordering. DNA containing internal biotin-TEG and internal etheno-dA requires an off-catalog order.

**Table S2.** Linear dsDNA prepared for this study. Below is a table of linear dsDNA templates that were prepared for this study, including the primers and template oligos used, DNA modifications, the PCR polymerase used, whether translesion synthesis was performed, how the resulting DNA was purified, and the figures in which each DNA template was used. The exact procedures used for DNA template preparation are described in Strobel, 2021 (16).

| ID | Fwd Primer | Rev Primer | Template Oligo | Modifications | PCR Polymerase | Translesion Synthesis | Clean Up | Used in Fig(s) |
| --- | --- | --- | --- | --- | --- | --- | --- | --- |
| 1 | TECD007 | LEGI10 | LEGH67 | -13,-30 dU;<br>5' biotin-TEG | Q5U | N/A | Gel Extracted | 2B, 5B, 5C, 5D, 5E, 6A, 6C, 6D |
| 2 | TECD007 | TECD001 | LEGH67 | -13,-30 dU;<br>Int biotin-TEG | Q5U | Yes | Gel Extracted | 3B, 3C, 5C, 5D, 5E, 6A, 6B, 6C, 6D, S2 |
| 3 | TECD007 | TECD000 | LEGH67 | -13,-30 dU;<br>Int desthiobiotin-TEG | Q5U | Yes | Gel Extracted | 3D |
| 4 | TECD009 | TECD001 | LEGH67 | 5' PC biotin;<br>Int C3 spacer;<br>-13,-30 dU;<br>Int biotin-TEG | Q5U | Yes | Thermolabile Exol<br>+ PCR clean-up | 4B, 4C, 4D. S1A |
| 5 | TECD006 | TECD002 | TECD016 | Int etheno-dA | Q5 | Yes | Thermolabile Exol<br>+ PCR clean-up | 4B; Note: this is the competitor DNA used for TEC purification |

**Table S3.**
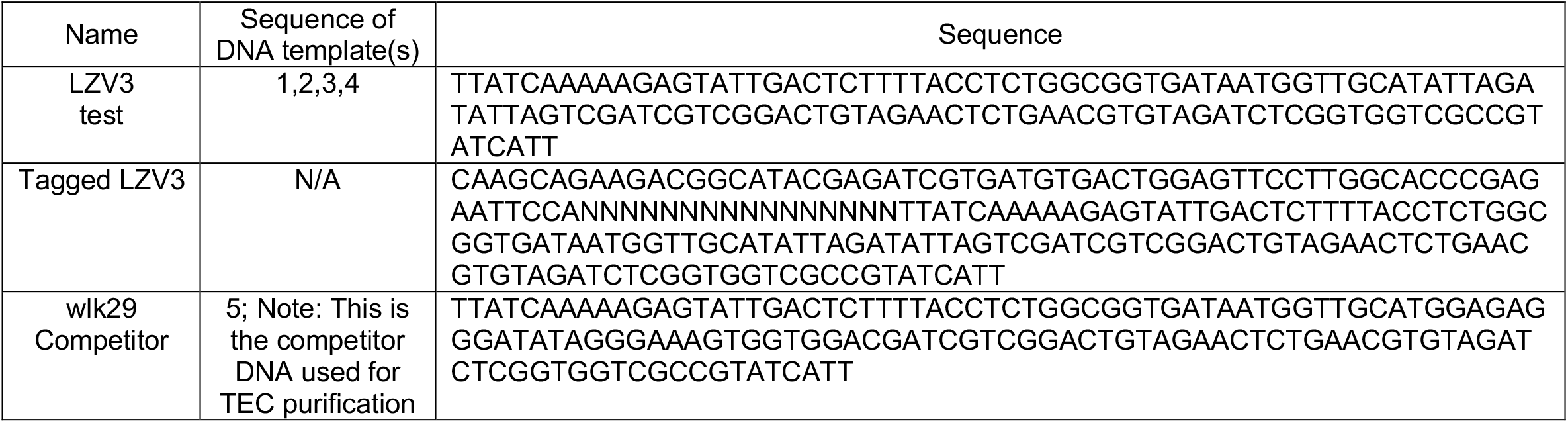
Linear dsDNA sequences. Below is a table of containing the sequence of each untagged and tagged linear dsDNA template used in this work. Fully annotated versions of LZV3 and Tagged LZV3 are available at Benching (See ‘Sequences’ in Materials and Methods for hyperlinks).

## References

1. Hug, H. and Schuler, R. (2003) Measurement of the number of molecules of a single mRNA species in a complex mRNA preparation. J Theor Biol, 221, 615–624.

2. Fu, G.K., Hu, J., Wang, P.H. and Fodor, S.P. (2011) Counting individual DNA molecules by the stochastic attachment of diverse labels. Proc Natl Acad Sci U S A, 108, 9026–9031.

3. Kivioja, T., Vaharautio, A., Karlsson, K., Bonke, M., Enge, M., Linnarsson, S. and Taipale, J. (2011) Counting absolute numbers of molecules using unique molecular identifiers. Nat Methods, 9, 72–74.

4. Nisson, P.E., Rashtchian, A. and Watkins, P.C. (1991) Rapid and efficient cloning of Alu-PCR products using uracil DNA glycosylase. PCR Methods Appl, 1, 120–123.

5. Nour-Eldin, H.H., Hansen, B.G., Norholm, M.H., Jensen, J.K. and Halkier, B.A. (2006) Advancing uracil-excision based cloning towards an ideal technique for cloning PCR fragments. Nucleic Acids Res, 34, e122.

6. Bitinaite, J., Rubino, M., Varma, K.H., Schildkraut, I., Vaisvila, R. and Vaiskunaite, R. (2007) USER friendly DNA engineering and cloning method by uracil excision. Nucleic Acids Res, 35, 1992–2002.

7. Geu-Flores, F., Nour-Eldin, H.H., Nielsen, M.T. and Halkier, B.A. (2007) USER fusion: a rapid and efficient method for simultaneous fusion and cloning of multiple PCR products. Nucleic Acids Res, 35, e55.

8. Bitinaite, J. and Nichols, N.M. (2009) DNA cloning and engineering by uracil excision. Curr Protoc Mol Biol, Chapter 3, Unit 3 21.

9. Vaisvila, R. and Bitinaite, J. (2013) Gene synthesis by assembly of deoxyuridine-containing oligonucleotides. Methods Mol Biol, 978, 165–171.

10. Lindahl, T., Ljungquist, S., Siegert, W., Nyberg, B. and Sperens, B. (1977) DNA N-glycosidases: properties of uracil-DNA glycosidase from Escherichia coli. J Biol Chem, 252, 3286–3294.

11. Lindahl, T. (1982) DNA repair enzymes. Annu Rev Biochem, 51, 61–87.

12. Dizdaroglu, M., Laval, J. and Boiteux, S. (1993) Substrate specificity of the Escherichia coli endonuclease III: excision of thymine- and cytosine-derived lesions in DNA produced by radiation-generated free radicals. Biochemistry, 32, 12105–12111.

13. Hatahet, Z., Kow, Y.W., Purmal, A.A., Cunningham, R.P. and Wallace, S.S. (1994) New substrates for old enzymes. 5-Hydroxy-2′-deoxycytidine and 5-hydroxy-2′-deoxyuridine are substrates for Escherichia coli endonuclease III and formamidopyrimidine DNA N-glycosylase, while 5-hydroxy-2′-deoxyuridine is a substrate for uracil DNA N-glycosylase. J Biol Chem, 269, 18814–18820.

14. Melamede, R.J., Hatahet, Z., Kow, Y.W., Ide, H. and Wallace, S.S. (1994) Isolation and characterization of endonuclease VIII from Escherichia coli. Biochemistry, 33, 1255–1264.

15. Jiang, D., Hatahet, Z., Melamede, R.J., Kow, Y.W. and Wallace, S.S. (1997) Characterization of Escherichia coli endonuclease VIII. J Biol Chem, 272, 32230–32239.

16. Strobel, E.J. (2021) Preparation of E. coli RNA polymerase transcription elongation complexes by selective photo-elution from magnetic beads. J Biol Chem, 100812.

17. Kwak, H., Fuda, N.J., Core, L.J. and Lis, J.T. (2013) Precise maps of RNA polymerase reveal how promoters direct initiation and pausing. Science, 339, 950–953.

18. Mahat, D.B., Kwak, H., Booth, G.T., Jonkers, I.H., Danko, C.G., Patel, R.K., Waters, C.T., Munson, K., Core, L.J. and Lis, J.T. (2016) Base-pair-resolution genome-wide mapping of active RNA polymerases using precision nuclear run-on (PRO-seq). Nat Protoc, 11, 1455–1476.

19. Campbell, E.A., Muzzin, O., Chlenov, M., Sun, J.L., Olson, C.A., Weinman, O., Trester-Zedlitz, M.L. and Darst, S.A. (2002) Structure of the bacterial RNA polymerase promoter specificity sigma subunit. Mol Cell, 9, 527–539.

20. Murakami, K.S., Masuda, S., Campbell, E.A., Muzzin, O. and Darst, S.A. (2002) Structural basis of transcription initiation: an RNA polymerase holoenzyme-DNA complex. Science, 296, 1285–1290.

21. Murakami, K.S., Masuda, S. and Darst, S.A. (2002) Structural basis of transcription initiation: RNA polymerase holoenzyme at 4 A resolution. Science, 296, 1280–1284.

22. Bae, B., Feklistov, A., Lass-Napiorkowska, A., Landick, R. and Darst, S.A. (2015) Structure of a bacterial RNA polymerase holoenzyme open promoter complex. Elife, 4.

23. Strobel, E.J., Lis, J.T. and Lucks, J.B. (2020) Chemical roadblocking of DNA transcription for nascent RNA display. J Biol Chem, 295, 6401–6412.

24. Boudsocq, F., Iwai, S., Hanaoka, F. and Woodgate, R. (2001) Sulfolobus solfataricus P2 DNA polymerase IV (Dpo4): an archaeal DinB-like DNA polymerase with lesion-bypass properties akin to eukaryotic poleta. Nucleic Acids Res, 29, 4607–4616.

25. Fiala, K.A., Brown, J.A., Ling, H., Kshetry, A.K., Zhang, J., Taylor, J.S., Yang, W. and Suo, Z. (2007) Mechanism of template-independent nucleotide incorporation catalyzed by a template-dependent DNA polymerase. J Mol Biol, 365, 590–602.

26. Olszewski, M., Rebala, K., Szczerkowska, Z. and Kur, J. (2005) Application of SSB-like protein from Thermus aquaticus in multiplex PCR of human Y-STR markers identification. Mol Cell Probes, 19, 203–205.

27. Sarkar, G., Kapelner, S. and Sommer, S.S. (1990) Formamide can dramatically improve the specificity of PCR. Nucleic Acids Res, 18, 7465.

28. Mattila, P., Korpela, J., Tenkanen, T. and Pitkanen, K. (1991) Fidelity of DNA synthesis by the Thermococcus litoralis DNA polymerase--an extremely heat stable enzyme with proofreading activity. Nucleic Acids Res, 19, 4967–4973.

29. Holz, K., Pavlic, A., Lietard, J. and Somoza, M.M. (2019) Specificity and Efficiency of the Uracil DNA Glycosylase-Mediated Strand Cleavage Surveyed on Large Sequence Libraries. Sci Rep, 9, 17822.

30. Kosaka, H., Hoseki, J., Nakagawa, N., Kuramitsu, S. and Masui, R. (2007) Crystal structure of family 5 uracil-DNA glycosylase bound to DNA. J Mol Biol, 373, 839–850.

31. Fromme, J.C. and Verdine, G.L. (2003) Structure of a trapped endonuclease III-DNA covalent intermediate. EMBO J, 22, 3461–3471.

32. Shultzaberger, R.K., Chen, Z., Lewis, K.A. and Schneider, T.D. (2007) Anatomy of Escherichia coli sigma70 promoters. Nucleic Acids Res, 35, 771–788.

